# Skeletal and Dental Development Preserve Evidence of Energetic Stress in the Moose of Isle Royale

**DOI:** 10.1101/831156

**Authors:** C. Brown, C.E. Rinaldi, W. J. Ripple, B. Van Valkenburgh

## Abstract

Food shortages can leave diagnostic, and in the case of the dentition, irreversible changes in mineralized tissue that persist into historical and fossil records. Consequently, developmental defects of tooth enamel might be used to track ungulate population irruption but dental tissue’s capacity for preserving historical population density changes has yet to be investigated in wild populations. We test the ability of enamel defects, mandible and metapodial lengths to track changes in the well-known fluctuating moose population of Isle Royale National Park. Our study demonstrates that 1) a moose density threshold exists on the island above which there is a significant decrease in mandible and metatarsus length and a concomitant increase in enamel hypoplasias, 2) food limitation has a more pronounced effect on male than female skeletal growth, and 3) combined data from tooth enamel hypoplasias and bone lengths reflect the relative density of this ungulate population, and should be broadly applicable to other ungulate osteological samples. Developmental defects in dental enamel were among the highest recorded in a wild population, and even during low-density intervals the population density of Isle Royale moose has been high enough to negatively impact skeletal and dental growth, indicating the comparatively poor health of this century-old ecosystem

## 1 Introduction

Mammalian teeth and skeletons contain a wealth of information about the life history of individuals and thus can preserve information about populations. Slowed or interrupted growth results from severe metabolic stress, such as might occur in association with an individual’s birth, weaning, episodes of disease, and poor nutrition. To test whether skeletal features track changes in population density, we chose ungulates, the prey base of most human societies and terrestrial large mammal carnivores for the past 50 million years. Ultimately, we hope to complement existing methods of estimating carnivoran(B. Van Valkenburgh and Hertel 2017) and proboscidean(Fisher 2001) population dynamics from skeletal material, allowing better characterization of Holocene and Anthropocene ecosystems. Such an approach requires skeletal characters that can be sampled non-destructively and have a high probability of surviving taphonomic processes. We identified such characters using skeletal material from a well-characterized, insular moose (*Alces alces*) population inhabiting Isle Royale National Park, Michigan.

Isle Royale, an approximately 544 km^2^ island in Lake Superior, was colonized by moose early in the twentieth century via a rare ice bridge. Isolated 90km off the shore of Ontario, the moose population is unique in that both immigration and emigration are negligible and there is only a single predator, the gray wolf (*Canis lupus*), that contributes to moose mortality. The lack of a complete predator guild and steep declines in the wolf population have allowed the moose population to fluctuate, punctuated by dramatic population crashes in 1934 and 1996 (Murie 1934), isleroyalewolf.org). Over the past 58 years, this isolated predator-prey system has been meticulously documented, facilitating assessment of the effects of climate, population size, and predation on ungulate health and mortality. Throughout this time period, Isle Royale’s lowest moose population density has exceeded that of nearby mainland areas, which often fall below 0.03 moose/km^2^ [IRNP minimum 0.71; mainland mean 0.55 +−.647] (Ripple and Beschta 2012). Resource limitation resulting from elevated densities has been associated with decreased carcass weight in multiple North American moose populations(Ferguson, Bisset, and Messier 2000), and is argued to be the mechanism driving the rapid dwarfing of the now century-old Isle Royale moose population (R. O. Peterson et al. 2011; Silvia et al. 1979).

Several studies have suggested that a variety of skeletal anomalies accompany poor juvenile nutrition in cervids. For example, delayed tooth emergence(R. O. Peterson, Schwartz, and Ballard 1983), shortened metatarsal bones(R. O. Peterson et al. 2011; Silvia et al. 1979; Geist 1998), osteoarthritis(R. O. Peterson et al. 2010), and osteoporosis(Hindelang and Peterson 1996) have all been linked to malnutrition. Reduced cranial size has been specifically associated with food shortages driven by elevated population density (Hoy, Peterson, and Vucetich 2018). Mandible length is of particular interest as the mandible has a high growth priority over other bones shortly after birth that slows only in the second year of life(Huot 1988). Prior to the 1996 Isle Royale population crash, shortened mandibles were thought to be linked to elevated population densities (R. O. Peterson and Holt 1996) as had been suggested for caribou(Reimers 1972). However, a study of female moose from another population (Ferguson, Bisset, and Messier 2000) found that density affects moose growth rate, not final size. In our study, we characterize the effect of density on tooth and bone growth in males and females, on and off the island.

Developmental defects of tooth enamel have been used to compare population health in two herds of caribou (Wu et al. 2012), but not yet to track shifts in ungulate population density over decades. The permanent dentition is a prime candidate for preserving evidence of irruptions because tooth tissue does not remodel. The crowns of moose permanent cheek teeth form over about a year, from roughly three months before birth to six to eight months post-weaning(R. L. Peterson 1950). Thus, malnutrition can affect the development of tooth enamel during this interval both directly through lack of forage, and indirectly, through an inadequately nourished mother. If foraging juveniles are insufficiently nourished, tooth development will be slowed and/or interrupted, leaving diagnostic features that persist into adulthood. In laboratory and domesticated animals, caloric restriction produces lines of arrested growth (LAGs) in bones, delayed tooth eruption, malocclusion, changes in mandibular shape, and dental enamel hypoplasias(Tonge 1965). Dental enamel hypoplasias (DEH) are among the most conspicuous, manifesting as a visible localized thinning or absence of enamel (Guatelli-Steinberg 2001; Goodman and Rose 1990). DEH are caused by impaired enamel secretion, a symptom of severe nutritional(Tonge 1965; Dobney and Ervynck 2000), pathological(Suckling, Elliott, and Thurley 1983; Goodman and Rose 1991, 1990; Hillson 1986) or environmental stress(Byerly 2007; Franz-Odendaal, Chinsamy, and Lee-Thorp 2004; Niven, Egeland, and Todd 2004; Wu et al. 2012). The morphology of DEH can vary with the degree of metabolic stress (Witzel et al. 2006), but DEH are easily identified in ungulates with exposed enamel surfaces, such as moose. In wild ungulates, hypoplasias have been attributed to weaning and seasonal stressors in extant and extinct giraffids (*Giraffa, Sivatherium*)(Franz-Odendaal 2004; Franz-Odendaal, Chinsamy, and Lee-Thorp 2004); bison (*Bison bison*)(Niven, Egeland, and Todd 2004; M. C. Wilson 1988; Byerly 2007; Barrón-Ortiz et al. 2019); horses (*Equus spp.*)(Barrón-Ortiz et al. 2019); white deer (*Odocoileus virginianus*(H. S. Davis 2013)); and caribou (*Rangifer tarandus groenlandicus*)(Wu et al. 2012).

To develop a method broadly applicable to archeological and paleontological samples, we focus on the size of two skeletal elements that are often preserved, mandibles with teeth and metapodials(O’Connor 2000). In addition, we quantified the frequency of growth anomalies (DEH) in the enamel of the cheek teeth. When compared with animals weaned in low-density years, moose that develop during periods of high population density were predicted to exhibit higher frequencies of anomalies in tissues that form after weaning (enamel, mandibular bone) in addition to shortened stature.

If combined dental and skeletal measures reliably track known changes in the moose population, and the development of these characters is conserved in ungulates, then we should be able to use them to identify relative density changes in other populations. We stress that this approach cannot be applied without acknowledging shifts in climate, forage species, and competitors. Even with approximate knowledge of these factors, assessments of nutritional stress using enamel surfaces and bone lengths are primarily a means of gauging population density relative to food availability, and not absolute levels of population density.

## Methods

### Materials

Skeletal specimens of moose collected from Isle Royale National Park (IRNP) between 1959 and 2019 are housed in the Michigan Technical Institute Mooseum in Alberta, Michigan. Our study sample spans 1959-2009. Samples collected prior to 1980 are primarily wolf kills whereas animals that died after that year were largely a result of overwinter mortality, the result of the wolf population being functionally extinct (J. and P. R. Vucetich 2012). Moose with associated mandibles and metatarsal bones were preferentially sampled. Sample sizes for the mandible, metatarsal, and dental analyses varied in size due to differential availability of specimens (see below). Most IRNP moose have been aged by Isle Royale Wolf Project (IRWP) scientists based on cementum annuli, allowing individuals to be assigned birth years. For 20 IRNP mandible specimens without cementum ages, we estimated a birth year interval based on the specimen’s autopsy number (indicative of date of death) and age class at death (e.g. old adult, prime adult). Moose mandible and metatarsus lengths, Isle Royale climate data, and moose density data were downloaded from the IRWP website(J. and P. R. Vucetich 2012). For comparison, we also collected dental defect and mandible size data from the closest mainland Ontario populations. The mainland sample (hereafter abbreviated as ONT) spanned the period 1950-1970, and thus experienced the same climate conditions as our earliest IRNP low density study period (1959-1963). ONT specimens are housed at the Royal Ontario Museum in Toronto, Canada. Population density of the ONT population was estimated as between 0.55-0.71 individuals per km^2^ using Canadian Department of Lands and Forests survey data and maps provided by the Canadian Ministry of Wildlife. The ONT moose population was subject to gray wolf, black bear (Ursus americanus) predation as well as human hunters, while the IRNP moose population had only gray wolves as predators. To assess the impact of population density on skeletal and dental characters, we analyzed data on mandible length (IRNP n=906; ONT n=36), metatarsus length (IRNP n=62), and enamel hypoplasias (IRNP n=138, ONT n=71). All dental specimens are listed in Supplementary Table 1.

### Population density threshold

The IRNP moose population has been unstable since the beginning of detailed study in 1959 (Figure 1). Previous work suggests that a density greater than two moose per km^2^ can result in stunted bone growth, particularly in the length of the mandible(R. O. Peterson and Holt 1996). In addition, the carrying capacity of the island, as determined by browse availability surveys made between 1945 and 1970, was estimated to be 1000 moose (or 1.83 moose/km^2^). We used the average of these two density estimates (2,1.83) as our hypothesized density threshold (1.92) beyond which moose were expected to show evidence of stress. Using year of birth to assign moose to high or low density categories, there was a significant correlation between population density and skeletal/dental characters. There is evidence that overpopulation and resource limitation can impact moose growth during the year they are born, or during gestation(Palsson 1952; Solberg et al. 2004) or perhaps even earlier because ungulate population growth has been linked to conditions two years prior to birth (Patterson and Power 2002; Post and Stenseth 1998). We explored these other possible correlations to assess which years of high density has the greatest effect on skeletal and dental growth. We searched for correlations between our skeletal measures, DEH, and three possible years (birth, gestation, two years prior to birth) to assign moose to high or low density intervals using the Maximal Information Coefficient (MIC). The MIC assesses both linear and non-linear relationships between variables(Reshef et al. 2011). Analysis was conducted in R 3.3.2 (R Core team) using the package Minerva(Albanese et al. 2013). The analysis was bootstrapped to obtain confidence intervals and p values for each correlation.

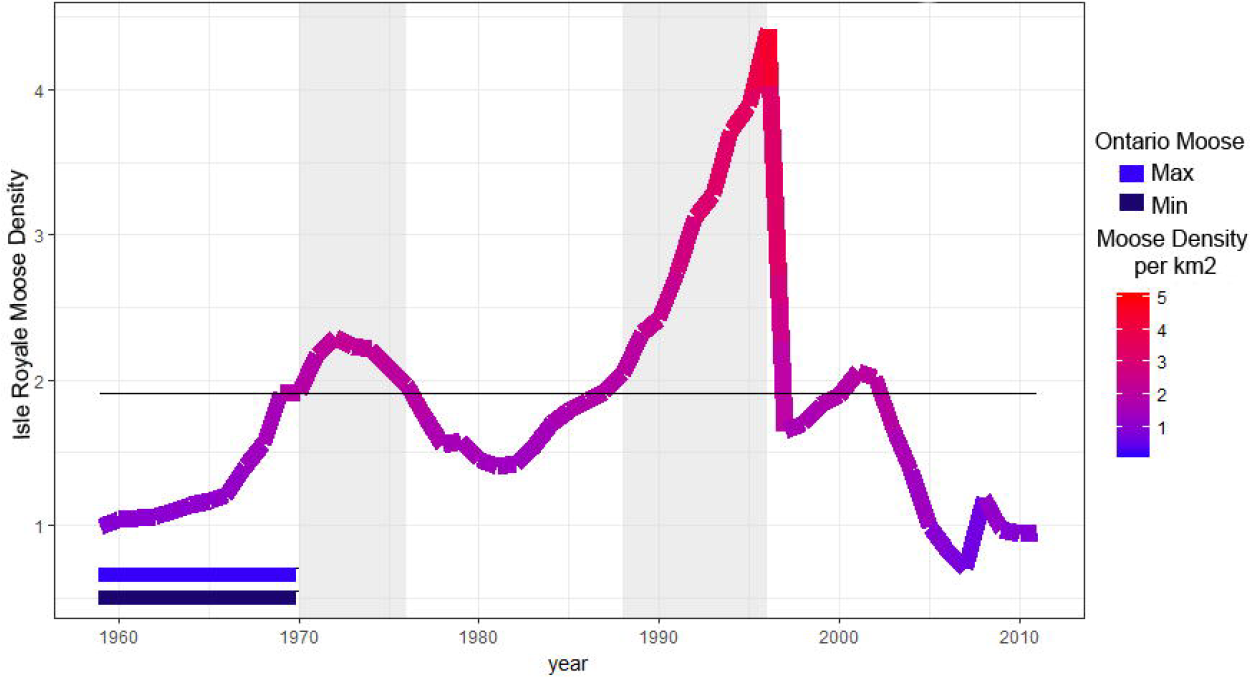

Based on the year with the strongest correlation (birth year, see below) the density threshold divided IRNP samples into five time periods: two of low density, two of high density and one transition period (Figure 1, Table 1). The most recent time period (1997-2009) is anomalous due to fluctuating population densities following a 75% reduction in population size. Moose from this time were born under densities ranging from 0.82-2.02 moose per km^2^ with an average density of 1.86, just below our density threshold. We could not subdivide this time period and retain enough samples for meaningful statistical analyses, so we draw most of our conclusions from the first four time periods.

**Table 1.** Time periods of study, population density, frequency of dental enamel hypoplasias (DEH), average score of hypoplasias and mean bone length measures for two moose localities. Sample sizes are indicated in parentheses. IRNP= Isle Royale National Park; ONT= closest mainland Ontario moose.

| Study Period | Years | Density Range (moose/km<sup>2</sup>) | Frequency of Moose Exhibiting DEH | Frequency Exhibiting Post-Weaning DEH | Mean DEH per Dentary | Mean Mandible Length (mm) | Mean Metatarsus Length (mm) |
| --- | --- | --- | --- | --- | --- | --- | --- |
| IRNP 1 | (1959 - 1968) | 1.05-1.91 | 33% (21) | 9.5% | 1.54 | 357.5±12.69 | 388±9.13 |
| IRNP 2 | (1969 - 1974) | 1.92-2.33 | 80.95% (21) | 42.86% | 3.95 | 356.7±8.37 | 377.9±10.63 |
| IRNP 3 | (1977 - 1986) | 1.41-1.86 | 58.82% (34) | 29.41% | 1.87 | 361.7±8.91 | 385.72±11.2 |
| IRNP 4 | (1987 - 1996) | 1.92-4.04 | 65.21% (23) | 34.78% | 2.19 | 347.8±17.3 | 364±25.48 |
| IRNP 5 | (1997 - 2003) | 1.63-2.05 | 74.28% (35) | 29.62% | 2.24 | NA | 373.92±16.3 |
| ONT | (1950<br>-<br>1970) | 0.16-0.80 | 36% (71) | 4.41% | 1.02 | 374.07±19.1 | NA |

### Life history and tooth development in moose

We reconstructed the development of tooth crowns in relation to life history events using published and new radiographs. Life history data were taken from Isle Royale Annual Reports (isleroyalewolf.org) as well as Peterson (1950, 1977). We radiographed the mandibles of juveniles of various ages (Supplementary Figure 1) to supplement radiographs from Peterson (1950). The interval of tooth crown development was estimated as beginning in the first month in which radio-opaque tissue was visible in the tooth crypt and ending when the full crown was visible to the level of root bifurcation. These ranges were plotted alongside major life history events (Figure 2) to assess which tissues developed near the times of birth and weaning. Differences in male and female moose development that contribute to size differences are also indicated as life history events.

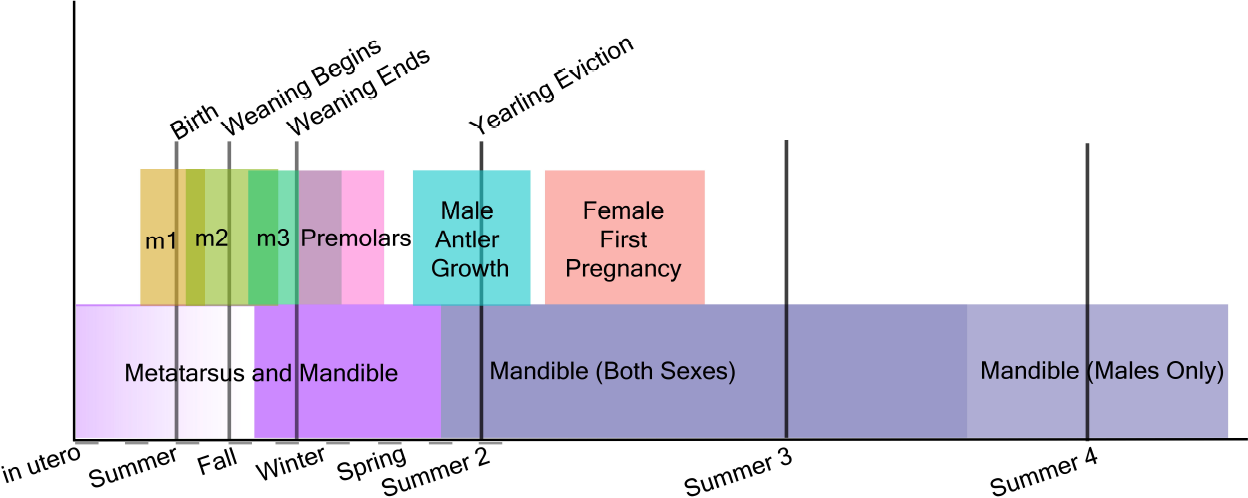

### Dental characters

All surveyed mandibles were radiographed in lateral view to reveal LAGs and root abnormalities. Two observers (CER,CB) independently surveyed 138 IRNP mandibles as well as 71 ONT mandibles (CB only) for six features: malocclusion, dental crowding, abnormal wear, tooth fracture, hypomineralization, and dental enamel hypoplasias (DEH). Of the six features, only DEH were observed. The widespread pitting, abnormal wear and characteristic chalky appearance of enamel affected by hyperflourosis(Hillson 1986; Shupe et al. 1987), another condition that causes enamel pitting in ungulates(H. Kierdorf et al. 2000; U. Kierdorf, Kierdorf, and Fejerskov 1993; U. Kierdorf et al. 1996) was absent.

DEH were categorized as pits, grooves, missing enamel, and linear enamel hypoplasias following the Fédération Dentaire Internationale Developmental Defects of Enamel Index ((Federation Dentarie Internationale 1982); Supplemental Figure 2). Hypoplasia morphology varies with stress severity(Witzel et al. 2006) and we sought to assess whether hypoplasias manifested differently over time on Isle Royale. Consequently, we assessed hypoplasia data in two ways, first, with all defects, and second, with only the more severe forms. Hypoplasia frequency distribution across groups was assessed using a bootstrapped chi-squared test. To analyze differences among time periods, we used Relative Risk Assessment. Commonly used in epidemiological studies (Simon 2001; McNutt 2003; Zhang and Yu 1998), Relative Risk Assessment estimates occurrence rates of relatively infrequent events and makes pairwise frequency comparisons between groups, such as the moose in each of our six samples (each of the five IRNP time periods plus the mainland).

### Mandible and metatarsus length

Mandible length was measured as the distance from the most ventral point of the mandibular notch to the most caudal point of the mandibular foramen (Supplemental Figure 3), permitting measurements on specimens with carnivore damage to the angle of the mandible. ONT mandible length data were captured by CB using digital photographs and ImageJ (http://imagej.nih.gov/ij/). IRNP adult mandibles less than 250mm in length (n=3) were considered outliers and excluded to facilitate meaningful statistical analysis (median adult length is 347.22 mm; median older adult length is 357.45mm).

Several factors can confound an assessment of the impact of population density on morphology, such as climate(J. A. Vucetich and Peterson 2010), moose sex, and age(Saether 1983). Climate has a significant effect on moose osteological development(Silvia et al. 1979; J. A. Vucetich and Peterson 2010) and the Isle Royale population’s growth rate(J. A. Vucetich and Peterson 2010); thus, it may be expected to play a role in the new characters assessed here. Moose sex and age can be expected to interact, as adult male moose continue to grow for several years past sexual maturity while females do not(Saether 1983). We analyzed bone lengths as pooled samples as well as subdivided into prime-age (<9 years) and old-age (9+ years) moose to avoid the confounding effect of age on male moose size at death. Because our length data do not follow assumptions of normality (mandible length, Shapiro-Wilkes W=0.79, p<2.2e−16, metatarsus length W = 0.89, p-value = 9.86e−07), we used a bootstrap in all analyses and chose tests robust to such data. To assess the effect of population density on moose mandible length between sexes and age groups we used a three-way ANOVA. To further analyze significant interactions between pairs of terms, such as sex and population density, we used two-way ANOVAs. For climate, we used the North Atlantic Oscillation (NAO) index, an established climate proxy for Isle Royale, and tested for correlations between the NAO and bone lengths, hypoplasia counts, and population density using the Maximal Information Coefficient (MIC) (Supplemental Figure 4). ONT specimens were not included in two- or three-way mandible analyses because many lacked sex data and were aged using the less reliable metric of tooth wear rather than cementum annuli (Pérez-barbería 2014). However, we did compare the ONT specimens to contemporaneous moose from IRNP period 1 using a bootstrapped two-group comparison. All code used in these analyses is available by request to CB. Dental data is available at Datadryad.org [https://doi.org/10.5068/D1237C]. IRNP mandible and metatarsus length data are property of the Isle Royale Wolf Project.

## Results

### Density threshold

Population density during a moose calf’s birth year produced the only significant MIC value for mandible lengths (MIC 0.134, p=0.002) and the strongest correlation with hypoplasia counts (MIC 0.198, p=0.017). Population density two years prior to birth was also significantly correlated with hypoplasia counts (MIC 0.157, p=0.009). MIC scores range from 0 (statistically independent) to 1 (no noise). For simplicity, we used the population density during an individual’s birth year to place them into groups of individuals that developed under high or low density conditions. Our results showed significant negative effects of birth-year density on mandible and metatarsus lengths and positive effects on the mean number of DEH per dentary (Table 1; see below).

### The effect of population size on teeth and stature

When all ages and sexes are pooled, IRNP moose born during the two high density periods exhibited significantly shorter mandibles (effect size = −4.89 mm, 95% CI [5.631, 10.418], p<0.0002) and metatarsals (effect size −6.097 mm, 95% CI [−5.943, 5.667], p<2e^−16^). They also exhibited elevated frequencies of dental anomalies (DEH), relative to moose born during the two low density periods (Table 1, Figures 3-4). Mandible length of IRNP moose varied only slightly over the first three time periods. It then declined sharply in period 4, the second of the high density periods. Metatarsal length also shows its steepest decline between periods 3 and 4 (Figure 4). The differences in size and DEH frequency over time suggest that overpopulation on IRNP had a negative impact on growth of teeth and bones. This is further supported by the comparison of mainland and IRNP moose. The frequency of DEH in IRNP moose ranges from 33% (period 1) to 81% (period 2) with a mean of 62.45% as opposed to 36% in the mainland moose sample. After period 1, the frequency of DEH in IRNP moose was 40%-200% greater than that observed in the ONT sample. Moose body size, measured by mandible length, was reduced on Isle Royale in the earliest period of study when compared to contemporaneous mainland moose. Although metatarsal length data was not available for the mainland sample, the moose residing on IRNP in period 1 had significantly shorter mandibles than the Ontario sample (effect size= 20.53mm, 95% CI [7.1155, −10.51], p=1e−04). However, the sample size of measurable Ontario mandibles was small (n=36) and skewed toward males and the sex ratio of the IRNP period 1 sample is skewed toward females (1.68:1) which probably accentuates the size difference.

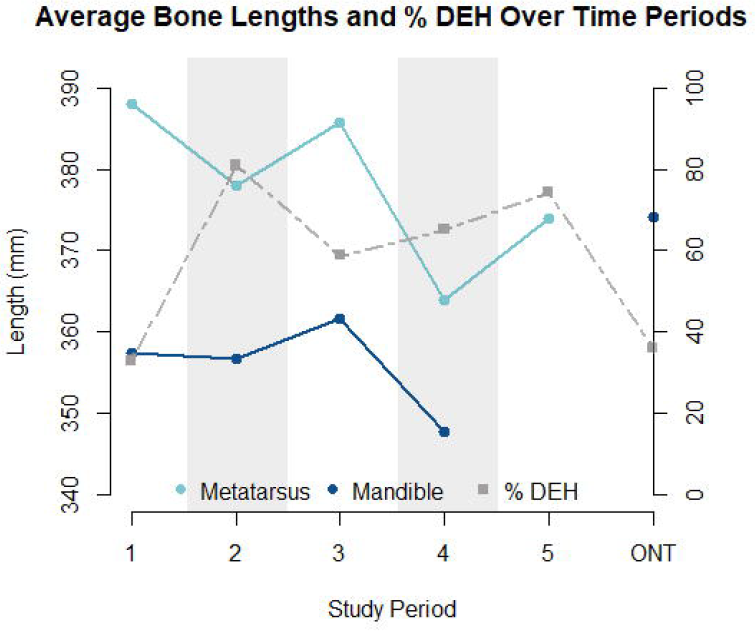

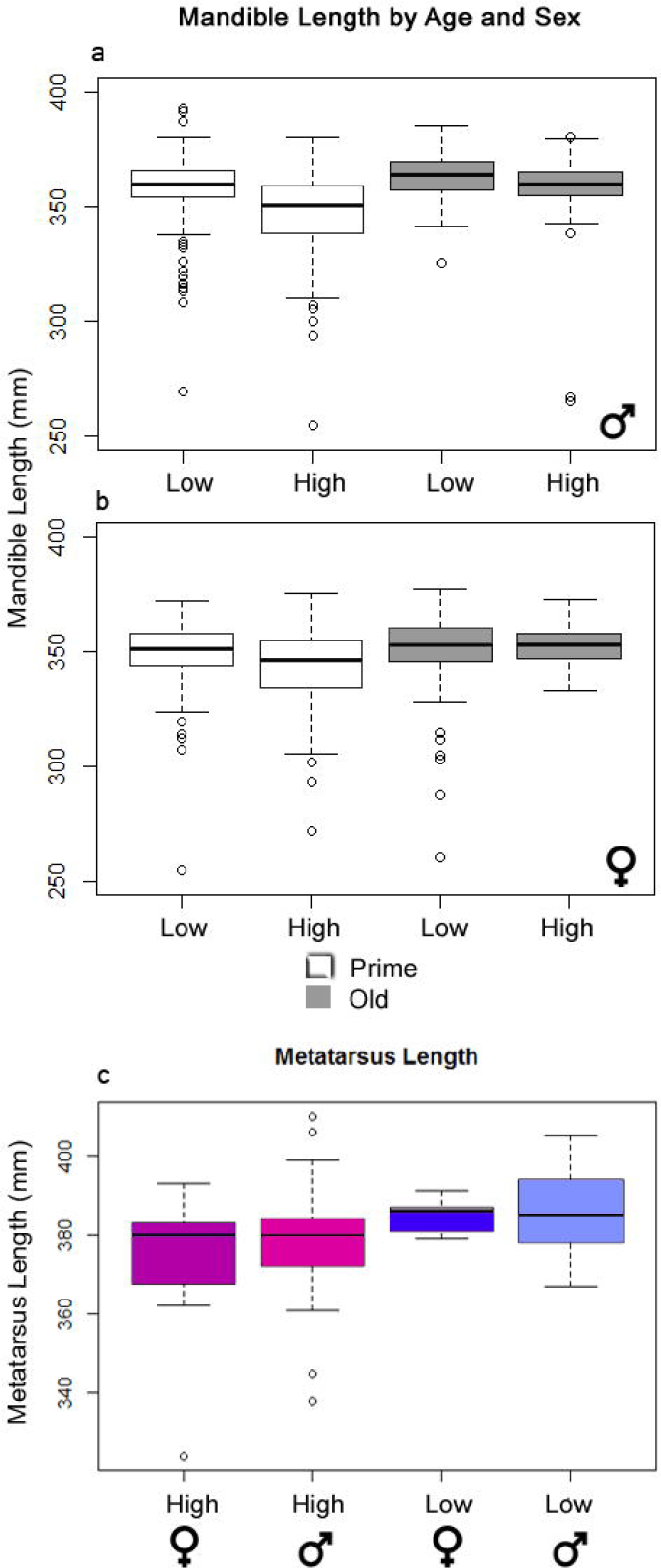

Individuals with shorter bone lengths tended to have more developmental enamel hypoplasias, and these were not evenly distributed across teeth and cusps (X^2^=473.37, df=5, n=418, p<.001). The premolar teeth are nearly free of hypoplasias whereas certain molar teeth, such as the m1 and m2, are particularly prone to them (Figure 5). Of all moose exhibiting enamel defects, 36.8% (77/209) developed a linear defect on the buccal surface of the m2 approximately 3 mm above the cemento-enamel junction while 22.9% (48/209) developed a linear defect on the buccal surface of m1. The number of post-weaning (estimated as after completion of m3) enamel hypoplasias per mandibular ramus was negatively correlated with mandible length (MIC 0.299, [0.275, 0.506], p=0.017), but mandible length and the total number of enamel hypoplasias were not (MIC 0.267, [0.267, 0.510], p=0.119). This implies that stressors other than post-weaning nutrition contribute to the total hypoplasia count. Likely candidates are some aspect of life history, such as birth or weaning (see Figure 2).

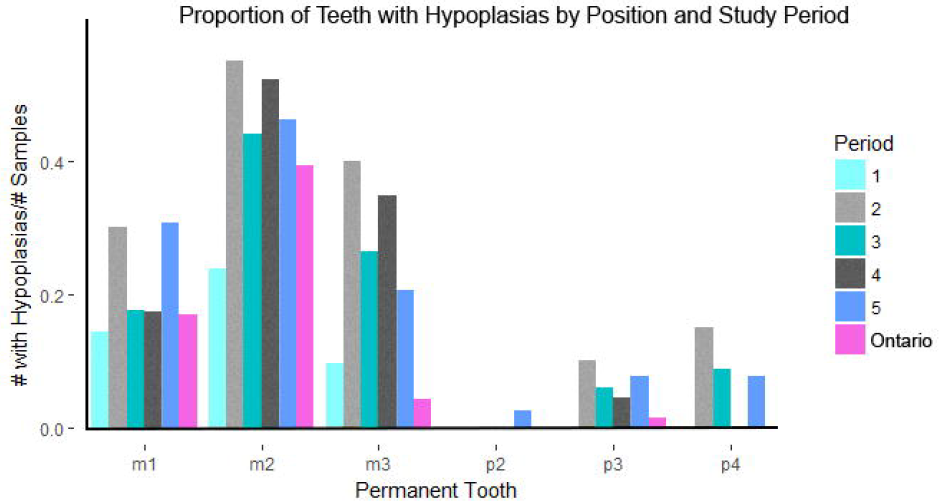

Dental enamel hypoplasias were recorded in all IRNP periods, but the frequency distribution is significantly uneven over time (X^2^=18.521, df=5, n=209, p<.001). DEH frequency peaks in period 2, declines in period 3, and then rises slowly across periods 4 and 5 (Figure 3). This pattern is driven by male moose; the proportion of females with enamel hypoplasias did not change significantly across periods (males X^2^= 9.5065, df = 4, n=45; p-value = 0.04961; females X^2^= 4.9811, df = 4, n=33, p-value = 0.2892). If premolars are formed entirely after weaning, then the relative risk of forming a defect increases once a moose stops nursing and forages. Risk of DEH in IRNP moose was also elevated during both high-density periods relative to low-density periods, and relative to the ONT sample (Table 2). IRNP moose from high-density period 2 had the greatest risk of forming DEH, particularly post-weaning hypoplasias that can be directly linked to insufficient forage. However, the risk of more severe pit and missing enamel DEH morphologies was not significantly elevated in any period (Table 3), demonstrating that while presence of DEH was associated with population density, DEH morphology was not.

**Table 2.** Relative Risk Assessment of incurring one or more dental enamel hypoplasias (DEH) on all teeth as well as teeth formed after weaning across time and localities. The relative risk of pit and missing enamel DEH reflects the occurrence of more severe and longer-lasting stress than the relative risk of all DEH types. IRNP= Isle Royale National Park; ONT= closest mainland Ontario moose.

| Period | vs. | Period | X times the risk of 1+ DEH (all morphologies) | X times the risk of 1+ post-weaning DEH (all morphologies) | X times the risk of pit or missing enamel morphology |
| --- | --- | --- | --- | --- | --- |
| 1 | vs. | 2 | .5797 | .281 | .3846 |
| 1 | vs. | 3 | .7173 | .425 | .5026 |
| 1 | vs. | 4 | .6521 | .359 | .5952 |
| 1 | vs. | 5 | .6366 | .609 | .7936 |
| 1 | vs. | ONT | 1.187 | 2.958 | 1.3 |
| 2 | vs. | 3 | 1.237 | 1.511 | 1.307 |
| 2 | vs. | 4 | 1.125 | 1.277 | 1.547 |
| 2 | vs. | 5 | 1.098 | 2.166 | 2.063 |
| 2 | vs. | ONT | 2.048 | 10.518 | 3.381 |
| 3 | vs. | 4 | .909 | .8455 | 1.184 |
| 3 | vs. | 5 | .887 | 1.434 | 1.579 |
| 3 | vs. | ONT | 1.65 | 6.96 | 2.587 |
| 4 | vs. | 5 | .976 | 1.69 | 1.333 |
| 4 | vs. | ONT | 1.820 | 8.23 | 2.184 |
| 5 | vs. | ONT | 1.864 | 4.854 | 1.368 |

**Table 3.**
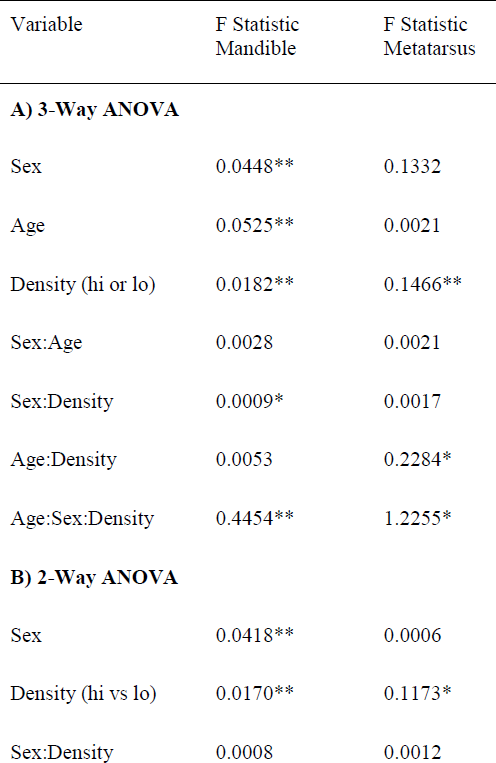
Bootstrapped ANOVA Results for various attributes. A single asterisk (*) indicates p<0.05 while a double asterisk (**) indicates p<0.01.

### Effect of age and sex of mandible and metatarsal lengths

Age, sex, population density, and the interaction between sex and population density all had a significant effect on the length of the mandible (Table 3). Pairwise investigations of sex, population density and bone length showed that high population density had a significant effect on median male mandible length (effect size= −5.28mm, 95% CI [−3.08, −9.33], p<0.001) but not median female mandible length (effect size =−0.55mm, 96%CI [−1.82, 2.31], p=0.59). As expected for a dimorphic species, male moose mandibles are significantly longer (Figure 4a,b); however the size difference is halved under high-density conditions (median difference at low density= 10.29mm; high density=5.33mm). Similarly, moose sex, population density and the interaction between sex and density were significant contributors to adult metatarsus length. Two-way analysis of metatarsus length, sex and density showed that population density, but not sex, had a significant association with metatarsal length on its own. The fact that sex alone was not significant is surprising given the mandible results and the overall size difference between males and females (Table 2, Figure 4) (mean difference 6.157cm, [−4.84, 16.521] p=0.5118). Although final metatarsus length is strongly influenced by *in utero* nutrition(Palsson 1952), adult metatarsus lengths were not significantly shorter in individuals with greater numbers of hypoplasias or hypoplasias incurred prior to weaning.

### Impact of climate

Climate, as estimated by our climate proxy, the North Atlantic Oscillation, was not strongly correlated with mandible length, metatarsus length, total DEH or post-weaning DEH (Supplementary Figure 4). While some MIC values were significant, the scatter of points was very large and the relationship detected by the MIC was both noisy and unclear when plotted.

## Discussion

Both dental enamel and bone lengths reflected long-term population trends in the Isle Royale moose population. Our analyses confirm that a moose density threshold exists on IRNP between 1.8 and 2 moose per km^2^, above which there is a significant decrease in mandible and metatarsus length and a concomitant increase in enamel hypoplasias. Density-dependent decreases in moose carcass weight, and presumably skeletal size, have been previously demonstrated(Ferguson, Bisset, and Messier 2000; Solberg et al. 2004). Here we show that signals of overpopulation may be preserved in both bones and teeth, raising the possibility of estimating relative population size from skeletal materials. The frequency of post-weaning enamel defects appears to reflect food-limitation. When sexes are pooled for the IRNP moose, the relative risk of post-weaning enamel defects is significantly higher under high-density conditions. There is a conspicuous near-absence of post-weaning hypoplasias in both low density, pre-crash (period 1) Isle Royale and mainland Ontario moose. Bone lengths are always significantly reduced on the island, but the effect is more pronounced when the population size is high. It is possible that wolf-killed moose samples may have skewed our IRNP samples toward smaller individuals. Nevertheless, we see a clear signal of resource limitation throughout time that reverses rapidly when conditions improve.

Comparison of moose on and off the island provided interesting insights into the historical health of Isle Royale. Our combined study of tooth and bone suggests that some Isle Royale individuals under low density conditions still experienced significantly more nutritional stress than their mainland counterparts, who experience densities five to ten times lower. Within Isle Royale NP, the lowest population densities are still much higher than is typical for moose, their mandible is shorter and DEH are more common. The earliest-studied Isle Royale moose had 1.18 times the chance of incurring a hypoplasia and 2.98 times the chance of incurring a post-weaning hypoplasia relative to their contemporaries on the mainland. Period 1 IRNP moose also had significantly shorter mandibles than the mainland moose and this might reflect the larger proportion of females in our IRNP period 1 sample. However, our results corroborate previous observations that moose began rapidly dwarfing within decades of their arrival on IRNP(R. O. Peterson et al. 2011). In addition to limited resources from their insular habitat, moose exerted heavy browsing pressure that altered forest composition through the suppression of woody plants(Murie 1934; Rotter and Rebertus 2015; Risenhoover and Maass 1987; McInnes et al. 1992) and extirpated important browse species(Murie 1934; Brandner, Peterson, and Risenhoover 1990; J. A. Vucetich and Peterson 2010). It takes more time for these species to recover than the periods that mark the cycling of the moose population(R. Peterson and McLaren 2010). In particular, localized shortages of the winter browse species balsam fir (*Abies balsamea*) may be linked to the prevalence of hypoplasias on late forming teeth in periods 2-5 (Figure 5), regardless of moose population density. Our results underscore the need for sustained periods of low moose population density to support both the moose and the vegetation on which they depend. This may be feasible given the National Parks Service’s recent decision to repopulate the island with wolves (Mlot 2018).

Signals preserved in bone and enamel demonstrate a clear effect of IRNP moose density at the population level and between the sexes. While the overall frequency of DEH was elevated under high density conditions, the risk of more severe pit and missing enamel DEH morphologies was not, indicating that overall DEH frequency is more informative than relative numbers of different types of DEH. DEH differed significantly in frequency between both sexes and population density regimes, with density-related changes driven almost entirely by males. Whereas Davis et. al (2013) found no difference in DEH frequency between the sexes in a sample of white-tailed deer (*Odocoileus virginianus*), we found that male moose had significantly elevated frequencies of DEH during high density. In contrast, female DEH frequency in IRNP moose was constant over time. Moose generation time is short(Geist 1963; Van Ballenberghe and Ballard 1994; Gaillard 2007) and male reproductive success is driven by body size(Garel et al. 2006). Consequently, male dental growth seems more affected by conditions early in life than females, due to the combined tendencies of males to grow faster and for a longer period of time(Kruuk et al. 1999; Solberg et al. 2004; Garel et al. 2006). In moose, it seems that the sexes differ in their sensitivity to metabolic stress as early as birth, the time at which the first permanent molar, the earliest-forming feature in our study, develops.

Across all five IRNP time periods and both the island and mainland populations, enamel hypoplasias appeared in constrained locations of the tooth crown that develop around the time of birth (m1) and weaning (m2). During high-density periods, there was a significantly greater risk of developing an enamel hypoplasia on tooth crowns formed both prior to and during weaning. This suggests that the development of DEH during weaning is most likely to occur in the presence of population-level resource limitation. This is similar to the phenomenon reported by Wu et. al. (2012) in which DEH associated putatively with birth were more common in a caribou population experiencing a population decline than non-declining populations.

The peak hypoplasia frequencies observed on Isle Royale (81%) are among the highest reported in wild populations of ungulates, with severe impairment of enamel formation visible on many individuals. A nutritionally stressed population of white-tailed deer exhibited defects in 27% of individuals(H. S. Davis 2013), while a population of caribou in decline had a frequency of 25.1% (Wu et al. 2012). Although a declining population of Alaskan moose exhibited higher levels of enamel defects (93%)(Stimmelmayr and Persons 2006), the defects were not localized lesions like the DEH described in IR moose and the above populations, but rather structural failures affecting most of the tooth crown that were putatively linked to local geochemistry(Clough, MacKenzie, and Broders 2010; MacKenzie et al. 2011). Isle Royale is also an unusual habitat give its insular nature and isolation for other populations. Given that IRNP moose do exhibit reduced heterozygosity(P. J. Wilson et al. 2003), it is possible, but unlikely, that their extremely high frequencies of DEH result from inbreeding. To date, DEH frequency has not been shown to increase in inbred populations; furthermore, known genetic enamel disorders are characterized by thinning and pitting of all enamel surfaces, abnormal tooth shape and malocclusion in humans(Wright, Johnson, and Fine 1993; Hillson 1986) and mouse models(Gibson et al. 2001; Caterina et al. 2002; Poulter et al. 2014). In our IRNP moose samples, DEH appeared on otherwise normal teeth with no signs of abnormal wear due to malocclusion or other factors.

Unlike moose teeth, which complete their growth within a year or two at most, bones have the potential to capture the effects of food limitation over a longer period of time. Moose mandible and metatarsus lengths varied with population density as expected from existing life-history data; higher population density reduces food availability and consequently moose stature. Like studies of moose populations elsewhere (Solberg et al. 2008; Ferguson, Bisset, and Messier 2000; Garel et al. 2006), we found that high population density has a much greater effect on male than female size. Studying exclusively females, Ferguson et. al (2000) documented shortened mandibles in growing Canadian moose, but final adult mandible sizes did not differ under high-density conditions. We were able to quantify mandible lengths in both sexes and found a significant reduction in males but not females consistent with Ferguson et al. (2000). The greater impact of population density on males may again be attributed to their longer period of growth. Just as we observed from the tooth enamel, male skeletal growth slows while they are stressed, whereas females may compensate through other means such as delayed breeding (Reimers 1972; Sand 1996; Keech et al. 2000; Testa 2004; Garel et al. 2006).

Our study of IRNP moose demonstrated population-level changes in morphology, but also that there can be discrepancies between tooth enamel and bone length data. Mandible and metatarsus length tracked population density more closely than did DEH frequency (Figures 3-4). Within high density periods, the maximum population density is matched by minimum average bone lengths. Unexpectedly, moose from period 2 had the most frequent DEH and the greatest risk of incurring a DEH suggesting an extended period of stress, but not the greatest reduction in bone lengths. The extremely high population densities of period 4 had the most reduced bone lengths, but not the most frequent DEH. These contradictions may result from the longer growth period of bones in contrast to teeth. Because mandibles and metatarsi continue to grow for several years after birth, a reduction in growth in earlier years might be reversed in later years if food becomes plentiful. Compensatory growth later in life has been demonstrated in cervids and other ungulate species(Luke, Tonge, and Reid 1981; Watkins 1991; Wairimu and Hudson 1993; Freetly, Ferrell, and Jenkins 2000; Betterly 2011) and could obscure the population density signal found with our approach. Unlike bone length, which varies significantly between individuals and even cohorts of individuals on a similar nutritional plane (J. Vucetich, pers. comm. 2015), teeth complete their growth at specific times and are relatively constrained in size. Consequently, tooth enamel and DEH frequency should provide a more reliable record of metabolic stresses early in life than either mandible or metatarsal lengths. The mismatch between peak hypoplasia frequency and peak population size may indicate that conditions in the first months of life were more taxing in period 2. Stressors during this period could have included energetic expenses not captured by the NAO index. Examples are parasites or locomotor difficulties resulting from a dense winter snow crust. There will always be factors that cannot be accounted for, but we can observe relative changes and increase our confidence by including both bones and teeth.

Given that we conducted this study with a goal of applying our findings to ungulate populations throughout the historical and paleontological record, the likelihood of preservation of characters is just as important as their ability to record episodic stress. In the fossil record, dental hypoplasia frequency, particularly during post-weaning, is likely to better preserve a record of historical overpopulation than mandible or metatarsal length given the material properties of the tissues. Skeletal measures tracked population irruptions with greater fidelity, however, teeth do not continue to grow or remodel and are often preserved intact. Therefore, skeletal measures should be considered in conjunction with dental development, particularly in the context of prehistorical applications. We stress that the measures presented here provide only relative estimates of population density. Future work will assess the impact of life history on susceptibility to DEH using additional cervid species. Nonetheless, if Isle Royale were to be viewed on the coarser time scale of a century, evidence of the island’s overpopulation would be preserved in the absence of major climate changes or island-wide extinction of winter browse. This gives us confidence that this same ecosystem dynamic could be recovered from the past.

The ability to draw evidence from multiple fossilized tissues when estimating relative population size may be particularly useful in reconstructing the terminal Pleistocene. Ungulate populations declined at differing rates in advance of range shifts and extinctions. Assuming that the frequency of dental enamel defects reflects relative population health in fossil ungulates, it might be possible to track trends in Pleistocene populations of ungulates over hundreds or thousands of years, as they approach their extinction. Many techniques and hypotheses have been tested on ungulates from this pivotal time (for a small sample see (Barrón-Ortiz et al. 2019; Feranec, Hadly, and Paytan 2009; Stuart et al. 2004; Guthrie 2003)). Analysis of DEH frequency in conjunction with climate proxies can indicate whether populations of a species might have declined due to bottom-up (e.g. climate) limitations, as opposed to top-down (e.g., hunting) pressure (see (Fisher 2001; Barrón-Ortiz et al. 2019)).

An increase in population-wide DEH will indicate overpopulation, whether from an increase in numbers or a contraction of resource pools. Some populations of ungulates experienced an increase in DEH following regional climate shifts in the terminal Pleistocene. In Pleistocene equids and Bison, Barrón-Ortiz et al. found that enamel hypoplasia frequency rose following both climate changes and changes in dental wear, and the trends varied by region. Other populations may show a pattern similar to our observations: a rise in DEH in the absence of climate and dietary changes. These may indicate a release from predation pressure instead, as we witnessed on Isle Royale following the functional extinction of wolves. Conversely a low frequency of DEH would indicate a stable population not exceeding carrying capacity. Moreover, examining the ungulate dental indicators alongside data on tooth wear and fracture in co-existing predator species(B. Van Valkenburgh 2016; B. Van Valkenburgh and Hertel 2017) has the potential to provide new insights into predator-prey dynamics of ancient ecosystems.

## 3 Author Contributions

B.V.V. and W.J..R. conceived of the project, B.V.V., C.B., and C.E.R. designed the collections research and collected data, C.B. executed statistical analysis, B.V.V. and C.B. wrote the paper. All authors gave final approval for publication.

## 5 Acknowledgments

Thank you to the Isle Royale Wolf Project John Vucetich and Rolf O. Peterson, Burton Lim, Susan Woodward, Jacqueline Miller, John Kohrn, Wendy Binder, Charlie Cobb, Mark Dallas for your advice, specimen access and training.

## Data Availability Statement

The dataset generated for this study can be found in the DataDryad repository [https://doi.org/10.5068/D1237C]. Additional publicly available data analyzed for this study can be found at isleroyalewolf.org [https://www.isleroyalewolf.org/sites/default/files/documents/Data_wolves_moose_Isle_Royale_une2019.xlsx]. The bone length datasets for this manuscript are not publicly available becausethey belong to the Isle Royale Wolf Project. Requests to access the datasets should be directed to JA Vucetich.

## References

Albanese, Davide, Michele Filosi, Roberto Visintainer, Samantha Riccadonna, Giuseppe Jurman, and Cesare Furlanello. 2013. “Minerva and Minepyl◻: A C Engine for the MINE Suite and Its R, Python and MATLAB Wrappers” 29 (3): 407–8. https://doi.org/10.1093/bioinformatics/bts707.

Ballenberghe, Victor Van, and Warren B. Ballard. 1994. “Limitation and Regulation of Moose Populations: The Role of Predation.” Canadian Journal of Zoology 72 (Macnab 1985): 2071–77. https://doi.org/10.1139/z94-277.

Barrón-Ortiz, Christina I., Christopher N. Jass, Raúl Barrón-Corvera, Jennifer Austen, and Jessica M. Theodor. 2019. “Enamel Hypoplasia and Dental Wear of North American Late Pleistocene Horses and Bison: An Assessment of Nutritionally Based Extinction Models.” Paleobiology. https://doi.org/10.1017/pab.2019.17.

Betterly, Benjamin W. 2011. “Compensatory Growth in Moose and Its Relationship to Sex, Longevity, Intraspecific Competition, and Senescence.” Michigan Technological University. https://digitalcommons.mtu.edu/etd-restricted/92/.

Brandner, T. A., R. O. Peterson, and K. L. Risenhoover. 1990. “Balsam Fir on Isle Royale: Effects of Moose Herbivory and Population Density.” Ecology. https://doi.org/10.2307/1940256.

Byerly, Ryan M. 2007. “Palaeopathology in Late Pleistocene and Early Holocene Central Plains Bison: Dental Enamel Hypoplasia, Fluoride Toxicosis and the Archaeological Record.” Journal of Archaeological Science. https://doi.org/10.1016/j.jas.2007.01.001.

Caterina, John J., Ziedonis Skobe, Joanne Shi, Yanli Ding, James P. Simmer, Henning Birkedal-Hansen, and John D. Bartlett. 2002. “Enamelysin (Matrix Metalloproteinase 20)-Deficient Mice Display an Amelogenesis Imperfecta Phenotype.” Journal of Biological Chemistry 277 (51): 49598–604. https://doi.org/10.1074/jbc.M209100200.

Clough, M J, C S K MacKenzie, and H G Broders. 2010. “The Spatial Variation of Extreme Tooth Breakage in an Herbivore and Potential Age Structure Effects.” Annales Zoologici Fennici 47 (4): 261–71. https://doi.org/10.5735/086.047.0404.

Davis, H S. 2013. “Enamel Hypoplasia as an Indicator of Nutritional Stress in Juvenile White-Tailed Deer.” Georgia Journal of Science 41 (2):95.

Dobney, Keith, and Anton Ervynck. 2000. “Interpreting Developmental Stress in Archaeological Pigs◻: The Chronology of Linear Enamel Hypoplasia,” 597–607. https://doi.org/10.1006/jasc.1999.0477.

Federation Dentarie Internationale. 1982. “A Epidemiological Index of Developmental Defects of Enamel (DDE Index).” Intenational Dental Journal 32: 159–67.

Feranec, Robert S., Elizabeth A. Hadly, and Adina Paytan. 2009. “Stable Isotopes Reveal Seasonal Competition for Resources between Late Pleistocene Bison (Bison) and Horse (Equus) from Rancho La Brea, Southern California.” Palaeogeography, Palaeoclimatology, Palaeoecology. https://doi.org/10.1016/j.palaeo.2008.10.005.

Ferguson, Steven H, Alan R Bisset, and François Messier. 2000. “The Influences of Density on Growth and Reproduction in Moose Alces Alces.” Wildlife Biology 6: 31–39.

Fisher, Daniel C. 2001. “Season of Death, Growth Rates, and Life History of North American Mammoths.” In Proceedings of the Internation Conference on Mammoth Site Studies. Publication in Anthropology., edited by D West, 22:122–35. University of Kansas.

Franz-Odendaal, Tamara A. 2004. “Enamel Hypoplasia Provides Insights into Early Systemic Stress in Wild and Captive Giraffes (Giraffa Camelopardalis).” Journal of Zoology 263 (2): 197–206. https://doi.org/10.1017/S0952836904005059.

Franz-Odendaal, Tamara A., A Chinsamy, and J Lee-Thorp. 2004. “High Prevalence of Enamel Hypoplasia in an Early Pliocene Giraffid (Sivatherium Hendeyi) from South Africa.” Journal of Vertebrate Paleontology 24 (1): 235–44. https://doi.org/10.1671/19.

Freetly, H. C., C. L. Ferrell, and T. G. Jenkins. 2000. “Timing of Realimentation of Mature Cows That Were Feed-Restricted during Pregnancy Influences Calf Birth Weights and Growth Rates.” Journal of Animal Science 78 (11): 2790–96.

Gaillard, Jean-Michel. 2007. “Are Moose Only a Large Deer?: Some Life History Considerations.” Alces 43: 1–11.

Garel, Mathieu, Erling Johan Solberg, Bernt Erik Saether, Ivar Herfindal, and Kjell Arild Hogda. 2006. “The Length of Growing Season and Adult Sex Ratio Affect Sexual Size Dimorphism in Moose.” Ecology 87 (3): 745–58. https://doi.org/10.1890/05-0584.

Geist, Valerius. 1963. “On the Behaviour of the North American Moose (Alces Alces Andersoni Peterson 1950) in British Columbia.” Behaviour 20 (3): 377–416. https://doi.org/10.1163/156853963X00095.

Geist, Valerius. 1998. Deer of the World: Their Evolution, Behaviour, and Ecology. 1st ed. Stackpole Books.

Gibson, Carolyn W., Zhi An Yuan, Bradford Hall, Glenn Longenecker, Enhong Chen, Tamizchelvi Thyagarajan, Taduru Sreenath, et al. 2001. “Amelogenin-Deficient Mice Display an Amelogenesis Imperfecta Phenotype.” Journal of Biological Chemistry 276 (34): 31871–75. https://doi.org/10.1074/jbc.M104624200.

Goodman, Alan H., and Jerome C. Rose. 1991. “Dental Enamel Hypoplasias as Indicators of Nutritional Status.” Advances in Dental Anthropology.

Goodman, Alan H., and Jerome C Rose. 1990. “Assessment of Systemic Physiological Perturbations from Dental Enamel Hypoplasias and Associated Histological Structures.” American Journal of Physical Anthropology 33 (11 S): 59–110. https://doi.org/10.1002/ajpa.1330330506.

Guatelli-Steinberg, Debbie. 2001. “What Can Developmental Defects of Enamel Reveal about Physiological Stress in Nonhuman Primates?” Evolutionary Anthropology 10 (4): 138–51. https://doi.org/10.1002/evan.1027.

Guthrie, R Dale. 2003. “Rapid Body Size Decline in Alaskan Pleistocene Horses before Extinction.” Nature, 169–71. https://doi.org/10.1038/nature02070.1.

Hillson, Simon. 1986. Teeth. Cambridge Manuals in Archaeology.

Hindelang, Mary, and Rolf O. Peterson. 1996. “Osteoporotic Skull Lesions in Moose at Isle Royale National Park.” Journal of Wildlife Diseases. https://doi.org/10.7589/0090-3558-32.1.105.

Hoy, Sarah R., Rolf O. Peterson, and John A. Vucetich. 2018. “Climate Warming Is Associated with Smaller Body Size and Shorter Lifespans in Moose near Their Southern Range Limit.” Global Change Biology 24 (6): 2488–97. https://doi.org/10.1111/gcb.14015.

Huot, Jean. 1988. “Review of Methods for Evaluating the Physical Condition of Wild Ungulates in Northern Environments.” Québec: Centre d’études Nordiques 50.

Keech, Mark A., Terry R. Bowyer, Jay M. Ver Hoef, Rodney D. Boertje, Bruce W. Dale, and Thomas R. Stephenson. 2000. “Life-History Consequences of Maternal Condition in Alaskan Moose.” The Journal of Wildlife Management 64 (2): 450–62. https://doi.org/Doi10.2307/3803243.

Kierdorf, Horst, Uwe Kierdorf, Alan Richards, and Frantisek Sedlacek. 2000. “Disturbed Enamel Formation in Wild Boars (Sus Scrofa L.) from Fluoride Polluted Areas in Central Europe.” Anatomical Record 259 (1): 12–24. https://doi.org/10.1002/(SICI)1097-0185(20000501)259:1<12::AID-AR2>3.0.CO;2-6.

Kierdorf, U., H. Kierdorf, and O. Fejerskov. 1993. “Fluoride-Induced Developmental Changes in Enamel and Dentine of European Roe Deer (Capreolus Capreolus L.) as a Result of Environmental Pollution.” Archives of Oral Biology 38 (12): 1071–81. https://doi.org/10.1016/0003-9969(93)90169-M.

Kierdorf, U, H Kierdorf, F Sedlacek, and O Fejerskov. 1996. “Structural Changes in Fluorosed Dental Enamel of Red Deer (Cervus Elaphus L.) from a Region with Severe Environmental Pollution by Fluorides.” Journal of Anatomy 188 (Pt 1: 183–95. http://www.pubmedcentral.nih.gov/articlerender.fcgi?artid=1167646&tool=pmcentrez&rendertype=abstract.

Kruuk, L., T. H. Clutton-Brock, K. E. Rose, and F. E. Guinness. 1999. “Early Determinants of Lifetime Reproductive Success Differ between the Sexes in Red Deer.” Proceedings of the Royal Society B: Biological Sciences 266 (June): 1655–61. https://doi.org/10.1098/rspb.1999.0828.

Luke, D. A., C. H. Tonge, and D. J. Reid. 1981. “Effects of Rehabilitation on the Jaws and Teeth of Protein-Deficient and Calorie-Deficient Pigs.” Cells Tissues Organs 110 (4): 299–305. https://doi.org/10.1159/000145440.

MacKenzie, Cynthia S Kendall, Michael J. Clough, Hugh G. Broders, and Mike Tubrett. 2011. “Chemical and Structural Composition of Atlantic Canadian Moose (Alces Alces) Incisors with Patterns of High Breakage.” Science of the Total Environment 409 (24): 5483–92. https://doi.org/10.1016/j.scitotenv.2011.08.066.

McInnes, Pamela F., Robert J. Naiman, John Pastor, and Yosef Cohen. 1992. “Effects of Moose Browsing on Vegetation and Litter of the Boreal Forest, Isle Royale, Michigan, USA.” Ecology 73 (6): 2059–75. https://doi.org/10.2307/1941455.

McNutt, L.-A. 2003. “Estimating the Relative Risk in Cohort Studies and Clinical Trials of Common Outcomes.” American Journal of Epidemiology 157 (10): 940–43. https://doi.org/10.1093/aje/kwg074.

Mlot, Christine. 2018. “Classic Wolf-Moose Study to Be Restarted on Isle Royale.” Science 361 (6409): 1298–99.

Murie, Adolph. 1934. “Moose of Isle Royale.” University of Michigan Press 1 (25): 44.

Niven, Laura B, Charles P Egeland, and Lawrence C Todd. 2004. “An Inter-Site Comparison of Enamel Hypoplasia in Bison◻: Implications for Paleoecology and Modeling Late Plains Archaic Subsistence” 31: 1783–94. https://doi.org/10.1016/j.jas.2004.06.001.

O’Connor, Terry. 2000. “The Archaeology of Animal Bones.” American Anthropologist 104 (1): 206. http://eprints.whiterose.ac.uk/72766/.

Palsson, H. 1952. “Effects of the Plane of Nutrition on Growth and the Development of the Carcass Quality in Lambs Part I. The Effects of High and Low Planes of Nutrition at Different Ages.” The Journal of Agricultiural Science 42 (1-2): 1–92.

Patterson, Brent R., and Vince A. Power. 2002. “Contributions of Forage Competition, Harvest, and Climate Fluctuation to Changes in Population Growth of Northern White-Tailed Deer.” Oecologia 130 (1): 62–71. https://doi.org/10.1007/s004420100783.

Pérez-barbería, Francisco Javier. 2014. “Evaluation of Methods to Age Scottish Red Deer◻:” 294: 180–89. https://doi.org/10.1111/jzo.12166.

Peterson, Randolph L. 1950. North American Moose. University of Toronto Press

Peterson, RO, and B McLaren. 2010. “Wolves, Moose, and Tree Rings on Isle Royale.” Advancement Of Science 266 (5190): 1555–58.

Peterson, Rolf O, and Kathy Holt. 1996. “Annual Report of Wolves on Isle Royale.”

Peterson, Rolf O, A John, D Drummer, and Clark Spencer. 2010. “Ecology of Arthritis,” 1124–28. https://doi.org/10.1111/j.1461-0248.2010.01504.x.

Peterson, Rolf O, Charles C Schwartz, and Warren B Ballard. 1983. “Eruption Patterns of Selected Teeth in Three North American Moose Populations.” The Journal of Wildlife Management 47 (3): 884. https://doi.org/10.2307/3808633.

Peterson, Rolf O, John A Vucetich, Dean Beyer, and Mike Schrage. 2011. “Phenotypic Variation in Moose◻: The Island Rule and the Moose of Isle Royale” 47 (Lieberman 2009): 125–33.

Post, Eric, and Nils Chr Stenseth. 1998. “Large-Scale Climatic Fluctuation and Population Dynamics of Moose and White-Tailed Deer.” Journal of Animal Ecology 67 (4): 537–43. https://doi.org/10.1046/j.1365-2656.1998.00216.x.

Poulter, James A., Steven J. Brookes, Roger C. Shore, Claire E L Smith, Layal Abi farraj, Jennifer Kirkham, Inglehearn Chris F., and Alan J. Mighell. 2014. “A Missense Mutation in ITGB6 Causes Pitted Hypomineralized Amelogenesis Imperfecta.” Human Molecular Genetics 23 (8): 2189–97. https://doi.org/10.1093/hmg/ddt616.

Reimers, Eigil. 1972. “Growth in Domestic and Wild Reindeer in Norway.” The Journal of Wildlife Management. https://doi.org/10.2307/3799094.

Reshef, David N, Yakir A Reshef, Hilary K Finucane, Sharon R Grossman, Gilean Mcvean, Peter J Turnbaugh, Eric S Lander, Michael Mitzenmacher, and Pardis C Sabeti. 2011. “Detecting Novel Associations in Large Datasets.” Science 334 (6062): 1518–24. https://doi.org/10.1126/science.1205438.Detecting.

Ripple, William J, and Robert L Beschta. 2012. “Large Predators Limit Herbivore Densities in Northern Forest Ecosystems.” European Journal of Wildlife Research 58 (4): 233–47. https://doi.org/10.1007/s10344-012-0623-5.

Risenhoover, Kenneth L., and Steven A. Maass. 1987. “The Influence of Moose on the Composition and Structure of Isle Royale Forests.” Canadian Journal of Forest Research 17 (5): 357–64. https://doi.org/10.1139/x87-062.

Rotter, Michael C., and Alan J. Rebertus. 2015. “Plant Community Development of Isle Royale’s Moose-Spruce Savannas.” Botany 93 (2): 75–90. https://doi.org/10.1139/cjb-2014-0173.

Saether, Bernt-Erik. 1983. “Relationship between Mandible Length and Carcass Weight of Moose in Norway.” The Journal of Wildlife Management. https://doi.org/10.2307/3808199.

Sand, H. 1996. “Life History Patterns in Female Moose (Alces Alces): The Relationship between Age, Body Size, Fecundity and Environmental Conditions.” Oecologia 106 (2): 212–20. https://doi.org/10.1007/BF00328601.

Shupe, J. L., P. V. Christofferson, A. E. Olson, E. S. Allred, and R. L. Hurst. 1987. “Relationship of Cheek Tooth Abrasion to Fluoride-Induced Permanent Incisor Lesions in Livestock.” American Journal of Veterinary Research 48 (10): 1498–1503.

Silvia, William J, Rolf O Peterson, John A Vucetich, William F Silvia, and W Silvia. 1979. “Variation in Metatarsal Morphology Among Subgroups of North American Moose (Alces Alces).”

Simon, S D. 2001. “Understanding the Odds Ratio and the Relative Risk.” Journal of Andrology 22 (4): 533–36. https://doi.org/.

Solberg, Erling Johan, Mathieu Garel, Morten Heim, Vidar Grøtan, and Bernt-Erik Sæther. 2008. “Lack of Compensatory Body Growth in a High Performance Moose Alces Alces Population.” Oecologia 158 (3): 485–98. https://doi.org/10.1007/s00442-008-1158-z.

Solberg, Erling Johan, Anne Loison, Jean Michel Gaillard, and Morten Heim. 2004. “Lasting Effects of Conditions at Birth on Moose Body Mass.” Ecography 27 (5): 677–87. https://doi.org/10.1111/j.0906-7590.2004.03864.x.

Stimmelmayr, R, and K Persons. 2006. “Incisor Tooth Breakage, Enamel Defects, and Peridontitis in a Declining Alaskan Moose Population” 42.

Stuart, A. J., P. A. Kosintsev, T. F.G. Higham, and A. M. Lister. 2004. “Pleistocene to Holocene Extinction Dynamics in Giant Deer and Woolly Mammoth.” Nature. https://doi.org/10.1038/nature02890.

Suckling, Grace, D. C. Elliott, and D. C. Thurley. 1983. “The Production of Developmental Defects of Enamel in the Incisor Teeth of Penned Sheep Resulting from Induced Parasitism.” Archives of Oral Biology 28 (5): 393–99. https://doi.org/10.1016/0003-9969(83)90134-6.

Testa, J W. 2004. “Population Dynamics and Life History Trade-Offs of Moose (Alces Alces) in South-Central Alaska.” Ecology 85 (5): 1439–52. https://doi.org/10.1890/02-0671.

Tonge, H. 1965. “Severe Undernutrition in Growing and Adult Animals,” no. 14.

Valkenburgh, Blaire V A N. 2016. “Incidence of Tooth Breakage Among Large, Predatory Mammals Author (s): Blaire Van Valkenburgh Source◻: The American Naturalist, Vol. 131, No. 2 (Feb., 1988), Pp. 291–302 Published by◻: The University of Chicago Press for The American Society Of” 131 (2): 291–302.

Valkenburgh, Blaire Van, and Fritz Hertel. 2017. “Tough Times at La Brea◻: Tooth Breakage in Large Carnivores of the Late Pleistocene Author (s): Blaire Van Valkenburgh and Fritz Hertel Published by◻: American Association for the Advancement of Science Stable URL◻: Http://Www.Jstor.Org/Stable/2881931J” 261 (5120): 456–59.

Vucetich, JA and Peterson RO. 2012. “The Population Biology of Isle Royale Wolves and Moose: An Overview.” Isleroyalewolf.Org. 2012. https://isleroyalewolf.org/data/data/home.html.

Vucetich, John A, and Rolf O Peterson. 2010. “The Influence of Top-down, Bottom-up and Abiotic Factors on the Moose (Alces Alces) Population of Isle Royale.” Society 271 (1535): 183–89. https://doi.org/10.1098/rspb.2003.2589.

Wairimu, Saphida, and R. J. Hudson. 1993. “Foraging Dynamics of Wapiti Stags (Cervus Elaphus) during Compensatory Growth.” Applied Animal Behaviour Science 36 (1): 65–79. https://doi.org/10.1016/0168-1591(93)90099-B.

Watkins, Wg. 1991. “Compensatory Growth of Wapiti (Cervus Elaphus) on Aspen Parkland Ranges.” Canadian Journal of Zoology 69: 1682–88. http://www.nrcresearchpress.com/doi/abs/10.1139/z91-233.

Wilson, M.C. 1988. Bison Dentitions from the Henry Smith Site, Montana: Evidence for Seasonality and Paleoenvironments at an Avonlea Bison Kill. Edited by L.B. Davis. Saskatoon: Saskatchewan Archaeological Society.

Wilson, Paul J, Sonya Grewal, Art Rodgers, Rob Rempel, Jacques Saquet, Hank Hristienko, Frank Burrows, Rolf Peterson, and Bradley N White. 2003. “Genetic Variation and Population Structure of Moose (Alces Alces) at Neutral and Functional DNA Loci” 683: 670–83. https://doi.org/10.1139/Z03-030.

Witzel, Carsten, Uwe Kierdorf, Keith Dobney, Anton Ervynck, Sofie Vanpoucke, and Horst Kierdorf. 2006. “Reconstructing Impairment of Secretory Ameloblast Function in Porcine Teeth by Analysis of Morphological Alterations in Dental Enamel.” Journal of Anatomy 209 (1): 93–110. https://doi.org/10.1111/j.1469-7580.2006.00581.x.

Wright, J. T., L. B. Johnson, and J. D. Fine. 1993. “Developmental Defects of Enamel in Humans with Hereditary Epidermolysis Bullosa.” Archives of Oral Biology 38 (11): 945–55. https://doi.org/10.1016/0003-9969(93)90107-W.

Wu, Jessica P., Alasdair Veitch, Sylvia Checkley, Howard Dobson, and Susan J. Kutz. 2012. “Linear Enamel Hypoplasia in Caribou (Rangifer Tarandus Groenlandicus): A Potential Tool to Assess Population Health.” Wildlife Society Bulletin 36 (3): 554–60. https://doi.org/10.1002/wsb.175.

Zhang, Jun, and Kai F Yu. 1998. “What’s the Relative Risk?: A Method of Correcting the Odds Ratio in Cohort Studies of Common Outcomes\r10.1001/Jama.280.19.1690.” JAMA 280 (19): 1690–91.

