## Supplemental Table and Figures for "Skeletal and Dental Development Preserve Evidence of Energetic Stress in the Moose of Isle Royale"

Supplementary Material

### 1 Supplementary Figures and Tables

| Specimen No. | Institution | Birth Year | Sex |
| --- | --- | --- | --- |
| 136 | MTUM | 1959 | male |
| 215 | MTUM | 1959 | male |
| 229 | MTUM | 1959 | female |
| 332 | MTUM | 1959 | male |
| 333 | MTUM | 1962 | male |
| 334 | MTUM | 1962 | female |
| 336 | MTUM | 1962 | male |
| 349 | MTUM | 1961 | male |
| 427 | MTUM | 1962 | female |
| 452 | MTUM | 1962 | male |
| 460 | MTUM | 1960 | male |
| 504 | MTUM | 1962 | male |
| 506 | MTUM | 1961 | male |
| 555 | MTUM | 1962 | male |
| 557 | MTUM | 1962 | female |
| 564 | MTUM | 1963 | male |
| 566 | MTUM | 1963 | male |
| 600 | MTUM | 1961 | male |
| 663 | MTUM | 1962 | male |
| 680 | MTUM | 1962 | male |
| 873 | MTUM | 1970 | female |
| 1067 | MTUM | 1959 | male |
| 1071 | MTUM | 1971 | male |
| 1092 | MTUM | 1970 | male |
| 1193 | MTUM | 1973 | male |
| 1518 | MTUM | 1972 | male |
| 1526 | MTUM | 1972 | male |
| 1543 | MTUM | 1976 | male |
| 1548 | MTUM | 1970 | male |
| 1582 | MTUM | 1972 | male |
| 1753 | MTUM | 1972 | NA |
| 1827 | MTUM | 1972 | NA |
| 1850 | MTUM | 1972 | NA |
| 1854 | MTUM | 1972 | male |
| 1857 | MTUM | 1972 | male |
| 1880 | MTUM | 1972 | female |
| 1885 | MTUM | 1972 | male |
| 1902 | MTUM | 1972 | male |
| 1933 | MTUM | 1972 | female |
| 1960 | MTUM | 1972 | male |
| 1964 | MTUM | 1972 | male |
| 2003 | MTUM | 1972 | male |
| 2067 | MTUM | 1986 | male |
| 2070 | MTUM | 1986 | male |
| 2073 | MTUM | 1986 | male |
| 2084 | MTUM | 1977 | male |
| 2089 | MTUM | 1981 | male |
| 2097 | MTUM | 1985 | male |
| 2108 | MTUM | 1982 | male |
| 2109 | MTUM | 1979 | male |
| 2141 | MTUM | 1986 | male |
| 2142 | MTUM | 1977 | male |
| 2144 | MTUM | 1979 | male |
| 2163 | MTUM | 1978 | female |
| 2172 | MTUM | 1982 | male |
| 2185 | MTUM | 1979 | male |
| 2188 | MTUM | 1980 | male |
| 2228 | MTUM | 1978 | male |
| 2229 | MTUM | 1978 | male |
| 2230 | MTUM | 1982 | male |
| 2234 | MTUM | 1978 | male |
| 2247 | MTUM | 1982 | male |
| 2276 | MTUM | 1985 | male |
| 2282 | MTUM | 1979 | male |
| 2295 | MTUM | 1981 | male |
| 2299 | MTUM | 1985 | female |
| 2318 | MTUM | 1981 | male |
| 2319 | MTUM | 1981 | male |
| 2326 | MTUM | 1980 | male |
| 2327 | MTUM | 1980 | male |
| 2351 | MTUM | 1982 | male |
| 2370 | MTUM | 1981 | male |
| 2372 | MTUM | 1980 | male |
| 2427 | MTUM | 1983 | male |
| 3510 | MTUM | 1997 | male |
| 4046 | MTUM | 1993 | NA |
| 4048 | MTUM | 1993 | male |
| 4050 | MTUM | 1992 | male |
| 4054 | MTUM | 1993 | male |
| 4055 | MTUM | 1992 | male |
| 4057 | MTUM | 1993 | female |
| 4060 | MTUM | 1995 | male |
| 4066 | MTUM | 1993 | male |
| 4070 | MTUM | 1993 | female |
| 4072 | MTUM | 1986 | male |
| 4073 | MTUM | 1992 | male |
| 4074 | MTUM | 1989 | male |
| 4077 | MTUM | 1993 | male |
| 4080 | MTUM | 1990 | male |
| 4081 | MTUM | 1988 | female |
| 4085 | MTUM | 1992 | male |
| 4086 | MTUM | 1996 | male |
| 4087 | MTUM | 1989 | female |
| 4089 | MTUM | 1990 | male |
| 4093 | MTUM | 1989 | male |
| 4163 | MTUM | 1997 | NA |
| 4176 | MTUM | 1996 | male |
| 4204 | MTUM | 1992 | NA |
| 4205 | MTUM | 2003 | NA |
| 4237 | MTUM | 1998 | male |
| 4244 | MTUM | 1998 | male |
| 4259 | MTUM | 1996 | male |
| 4289 | MTUM | 1999 | NA |
| 4293 | MTUM | 1997 | female |
| 4310 | MTUM | 2001 | NA |
| 4320 | MTUM | 2000 | male |
| 4324 | MTUM | 2006 | female |
| 4329 | MTUM | 1997 | male |
| 4330 | MTUM | 1997 | male |
| 4333 | MTUM | 1998 | female |
| 4338 | MTUM | 1999 | male |
| 4348 | MTUM | 1997 | NA |
| 4356 | MTUM | 2000 | female |
| 4406 | MTUM | 1999 | male |
| 4410 | MTUM | 1997 | male |
| 4415 | MTUM | 1997 | male |
| 4419 | MTUM | 1997 | male |
| 4422 | MTUM | 1999 | male |
| 4424 | MTUM | 1997 | male |
| 4436 | MTUM | 1997 | NA |
| 4445 | MTUM | 1997 | male |
| 4446 | MTUM | 1997 | male |
| 4451 | MTUM | 1992 | male |
| 4454 | MTUM | 1998 | male |
| 4457 | MTUM | 1999 | male |
| 4459 | MTUM | 2001 | NA |
| 4467 | MTUM | 1993 | male |
| 4505 | MTUM | 1999 | male |
| 4529 | MTUM | 1997 | male |
| 4531 | MTUM | 1998 | female |
| 4534 | MTUM | 2000 | male |
| 4537 | MTUM | 1996 | male |
| 4548 | MTUM | 1996 | male |
| 4552 | MTUM | 2005 | male |
| 4565 | MTUM | 1999 | NA |
| 4570 | MTUM | 1999 | NA |
| 4575 | MTUM | 1999 | NA |
| 4615 | MTUM | 1991 | NA |
| 4649 | MTUM | 2002 | NA |
| 4686 | MTUM | 1999 | NA |
| 4706 | MTUM | 2003 | NA |
| 4766 | MTUM | 2005 | NA |
| 4959 | MTUM | 2003 | NA |
| 16387 | ROM | 1945 | male |
| 16683 | ROM | 1946 | male |
| 18399 | ROM | NA | male |
| 18413 | ROM | 1947 | female |
| 18445 | ROM | 1947 | male |
| 18451 | ROM | 1947 | male |
| 18471 | ROM | 1947 | male |
| 18512 | ROM | 1947 | male |
| 18516 | ROM | 1947 | male |
| 18847 | ROM | 1948 | female |
| 18850 | ROM | 1948 | male |
| 18851 | ROM | 1948 | male |
| 18860 | ROM | 1948 | female |
| 19293 | ROM | NA | male |
| 19294 | ROM | NA | male |
| 19329 | ROM | 1948 | male |
| 19338 | ROM | 1948 | male |
| 19341 | ROM | 1949 | male |
| 19356 | ROM | 1948 | male |
| 19366 | ROM | 1948 | male |
| 19367 | ROM | 1948 | male |
| 19460 | ROM | NA | male |
| 19491 | ROM | 1949 | male |
| 19516 | ROM | NA | male |
| 19524 | ROM | NA | male |
| 19551 | ROM | 1947 | male |
| 19601 | ROM | 1949 | male |
| 19912 | ROM | NA | male |
| 19912 | ROM | 1949 | male |
| 19957 | ROM | 1949 | male |
| 19973 | ROM | NA | female |
| 19990 | ROM | 1949 | male |
| 20195 | ROM | 1950 | male |
| 20327 | ROM | 1950 | female |
| 25910 | ROM | NA | NA |
| 25911 | ROM | 1953 | male |
| 25912 | ROM | NA | female |
| 25912 | ROM | 1953 | male |
| 25913 | ROM | 1953 | male |
| 25914 | ROM | 1953 | male |
| 25915 | ROM | 1953 | male |
| 25916 | ROM | NA | male |
| 25916 | ROM | 1953 | female |
| 25917 | ROM | NA | male |
| 25917 | ROM | 1953 | male |
| 25918 | ROM | NA | male |
| 25918 | ROM | 1953 | male |
| 25919 | ROM | NA | male |
| 25919 | ROM | 1953 | female |
| 25920 | ROM | 1953 | male |
| 33889 | ROM | 1951 | male |
| 101559 | ROM | 1953 | male |
| 101560 | ROM | NA | male |
| 101569 | ROM | 1952 | male |
| 101590 | ROM | NA | male |
| 101602 | ROM | NA | male |
| 101653 | ROM | NA | male |
| 101667 | ROM | 1951 | male |
| 101696 | ROM | NA | male |
| 101736 | ROM | NA | male |
| 101763 | ROM | 1953 | male |
| 101769 | ROM | NA | male |
| 101785 | ROM | NA | male |
| 101789 | ROM | NA | male |
| 101831 | ROM | 1951 | male |
| 101834 | ROM | NA | male |
| 115-53 | ROM | NA | NA |
| 126-53 | ROM | NA | NA |
| 14-23-25-1 | ROM | 1953 | NA |
| 239-53 | ROM | NA | NA |
| 99-53 | ROM | NA | NA |

**Supplementary Table 1.** Specimens assessed for tooth and jaw development, institutions and sexes. ROM= Royal Ontario Museum, Toronto, Canada. MTUM= Michigan Technological University Mooseum, Alberta, Michigan.

#### 1.1 Supplementary Figures

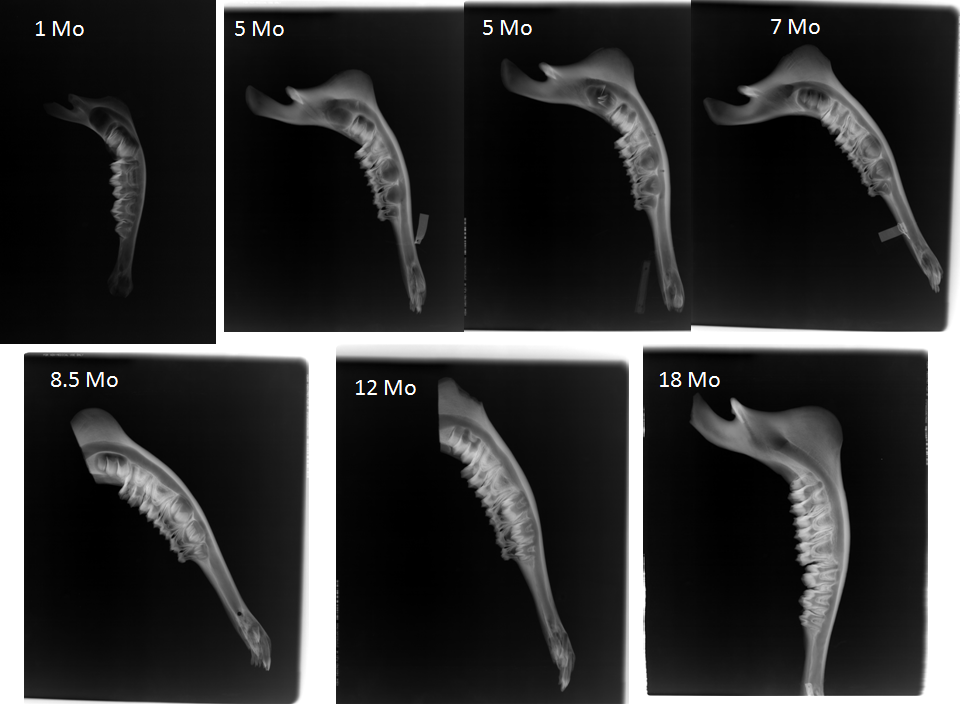

**Supplementary Figure 1.** Radiograph images of moose tooth development from birth to full adult dentition. Moose are born while the M1 crown is mineralizing; weaning occurs between 5 and 6 months of age while the M2 crown is mineralizing and the M3 is beginning to form. For additional radiographs, see Peterson (1950).

**Supplemental Figure 2.** Isle Royale dental enamel hypoplasia (DEH) morphologies, our assigned relative severity score, and a description of the ameloblast disruption that results in each morphology from Witzel et al. 2006.

| Hypoplastic Defect | Morphology | Score | Ameloblast Cohorts Affected |
| --- | --- | --- | --- |
| 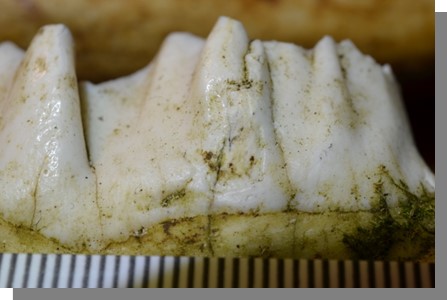 | Depression | 1 | Multiple; moderate impact and long duration |
| 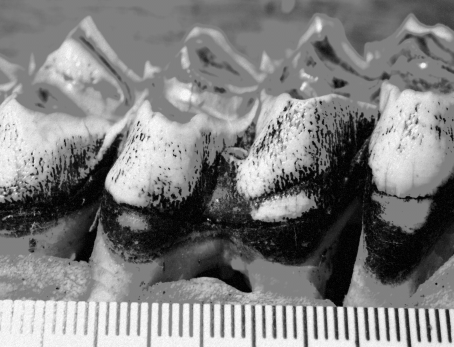 | Linear Enamel Hypoplasia (LEH), furrow morphology | 2 | Single; severe impact and very short duration |
| 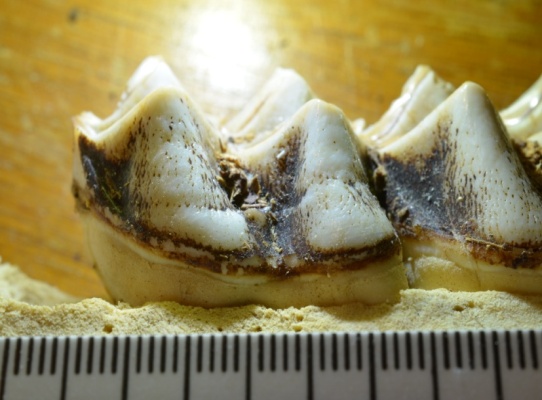 | Linear Enamel Hypoplasia (LEH), pit morphology | 2 | Single; severe impact and very short duration |
| 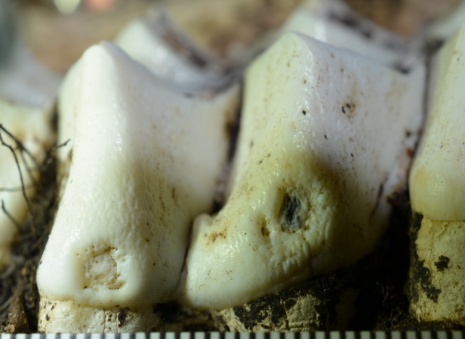 | Pit | 3 | Multiple; severe impact and short duration, thin enamel present on bottom of pit |
| 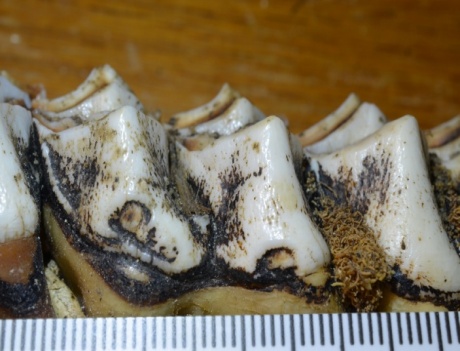 | Missing Enamel | 3 | Multiple; severe impact and short duration, enamel-secreting cells cease activity and tooth pulp (dentin) is exposed |

**Supplemental Figure 3.** Anatomical landmarks used for mandible length measurements. The solid yellow line represents measured jaw length. Image adapted from the Isle Royale Wolf Project.

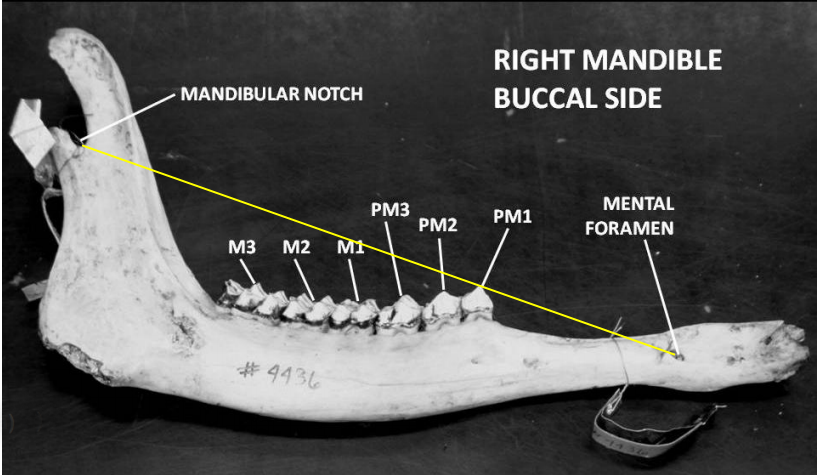

**Supplemental Figure 4.** The relationship between the North Atlantic Oscillation (NAO)--a proxy for winter severity--and surveyed skeletal and dental characters. The correlations between mandible length and population density during the birth year, gestation year and two years prior to birth were all weak and relatively noisy. A) NAO vs mandible length B) NAO vs total hypoplasia count C) NAO vs post-weaning hypoplasia count D) NAO vs metatarsus length.

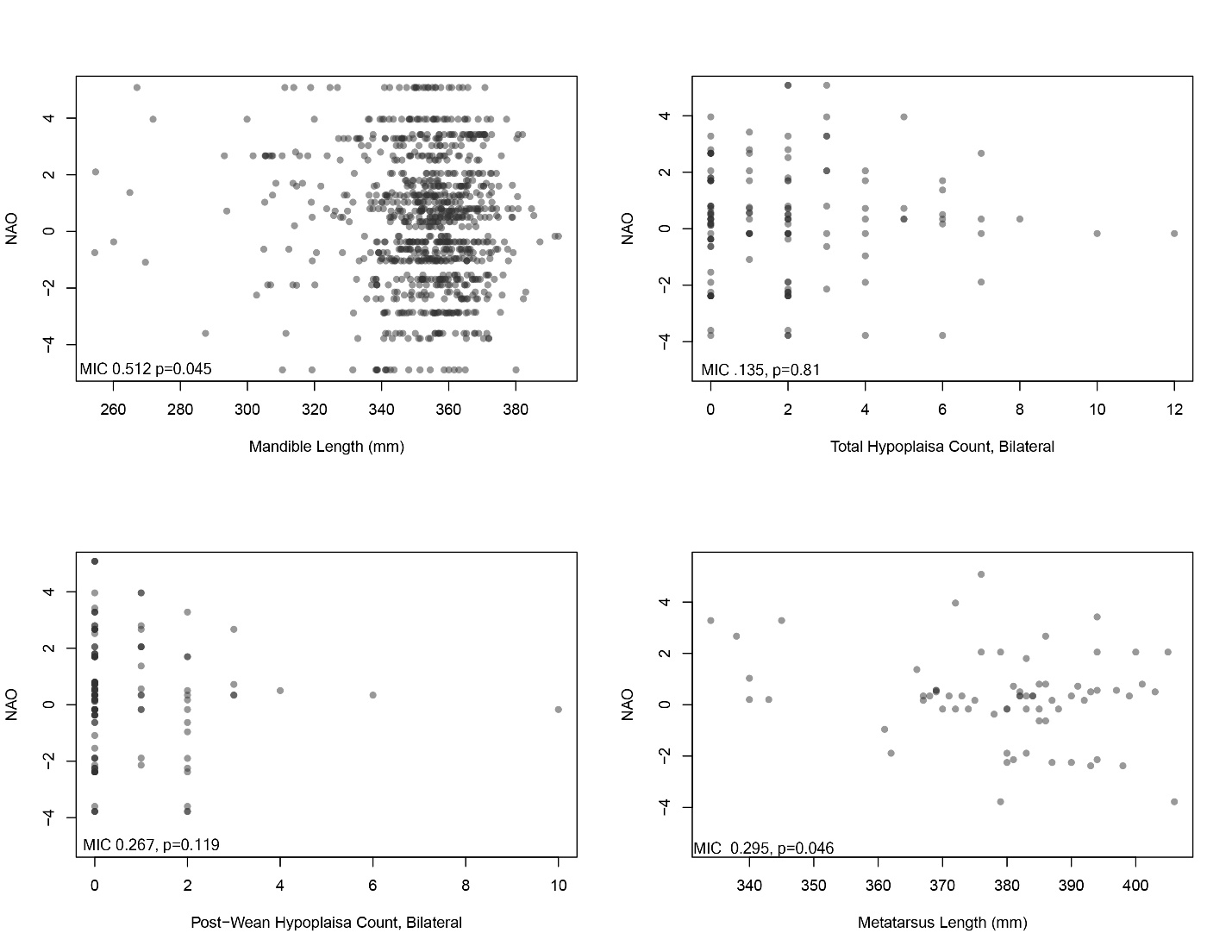

**Supplemental Figure 5.** Heatmap depiction of the relative risk of hypoplasia types between periods. Colors represent the risk of hypoplasia in the period on the y axis relative to the corresponding period on the x axis. Data plotted here can be found in table 2.3.

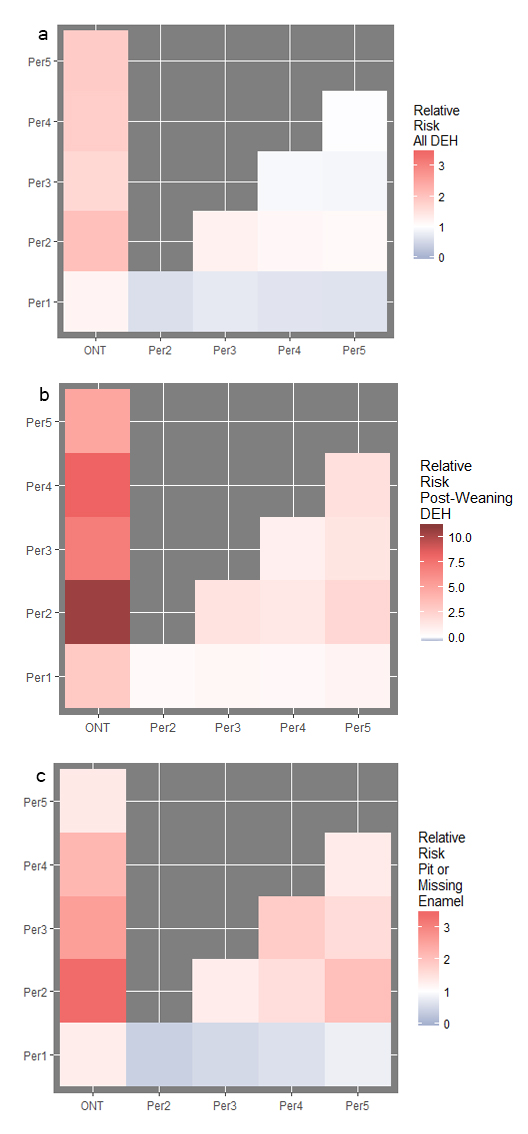
